# Neuraminidase-Mediated Desialylation Modulates Red Blood Cell Aggregation

**DOI:** 10.64898/2026.06.30.735505

**Authors:** Min Jin, Daria Tsvirkun, Chaouqi Misbah

**Affiliations:** Department of Endocrinology and Metabolic Diseases, The Third Affiliated Hospital of Sun Yat-sen Un; Universite Grenoble Alpes, CNRS, Laboratoire Interdisciplinaire de Physique, 38000, Grenoble, France

## Abstract

The glycocalyx of red blood cells (RBCs), a negatively charged surface layer rich in sialic acid residues, plays a crucial role in modulating RBC aggregation. In pathological conditions such as diabetes and sepsis, glycocalyx degradation is often observed along with abnormal RBC aggregation. However, the mechanistic relationship between these phenomena remains poorly defined. In this study, we investigate the effects of enzymatic glycocalyx degradation on RBC aggregation under physiologically relevant flow conditions. Using neuraminidase from *Clostridium perfringens* (*C. welchii*) at varying concentrations, we selectively removed sialic acid residues from the RBC glycocalyx, simulating different levels of desialylation observed in health and disease. Confocal microscopy confirmed the dose-dependent depletion of membrane sialic acid, while microfluidic experiments revealed a significant increase in both the size and stability of the RBC aggregates after enzymatic treatment. Our findings suggest that glycocalyx integrity is a crucial biophysical determinant of RBC aggregation, likely influencing both electrostatic repulsion and hydrodynamic forces. This study provides new insights into how the enzymatic modification of the glycocalyx contributes to pathological hemorheology and may inform future strategies for the diagnosis or treatment of vascular diseases.

## 1 Introduction

Red blood cells (RBCs) play a crucial role in oxygen delivery and microvascular perfusion. Their mechanical and adhesive properties are finely tuned to support reversible aggregation, especially under low shear conditions, and contribute to the non-Newtonian behavior of blood, balancing flow efficiency with effective capillary passage [1, 2, 3]. This behavior is essential for normal hemorheology and is largely influenced by the biophysical characteristics of the RBC membrane. A critical determinant of RBC aggregation lies in their surface properties, notably the glycocalyx, a dense layer composed of glycoproteins, glycolipids, and proteoglycans that confers a net negative charge through sialic acid residues [4, 5, 6]. Generally, glycocalyx helps intercellular recognition, communication, and intercellular adhesion [4]. This physiological negative charge provides electrostatic repulsion between neighboring cells, modulating interaction forces and preventing excessive aggregation.

Under physiological conditions, RBC aggregation is predominantly driven by plasma macromolecules such as fibrinogen, which bridge adjacent cells or induces osmotic depletion effects [2, 7, 8, 9]. These interactions are typically reversible and sensitive to shear stress. However, in certain pathological conditions, including diabetes, sepsis, and certain viral infections, RBC aggregation becomes markedly enhanced and sometimes irreversible [10, 11, 12, 13, 14]. While elevated levels of plasma macro-molecules have been linked to this phenomenon, alterations in the RBC glycocalyx are also frequently observed in these diseases [15, 16, 17, 18, 19], prompting the hypothesis that changes in membrane glycosylation may play a key role in altered RBC behavior. *In vivo* observations support the link between degraded RBC glycocalyx and enhanced RBC aggregation, for example, diabetic patients show elevated RBC aggregation under physiological flow (e.g. cephalic vein measurements) [20].

Among the notable changes in such pathological contexts is the degradation or modification of membrane-bound sialic acid residues. Neuraminidase (NA, also known as sialidase) is an enzyme that not only participates in physiological desialylation processes within mammalian cells [21] but is also commonly secreted by pathogens during infection [22]. By cleaving terminal sialic acids from glycoproteins and glycolipids, NA activity, which is often elevated in conditions such as influenza, diabetes, and sepsis [22, 23], diminishes the negative surface charge of RBCs and has been implicated in modulating intercellular adhesive forces and aggregation behavior [11, 23]. A key *in vitro* study further demonstrated that NA derived from *Clostridium perfringens* (*C. welchii*) efficiently removes sialic acid residues from RBC membranes, thereby altering surface marker expression and enhancing erythrophagocytosis [24].

Although previous studies have established links between glycocalyx degradation and altered RBC interactions, these investigations have largely been confined in static conditions or conventional bulk rheological measurements. Such approaches do not fully capture the dynamic behaviors of RBCs under physiologically relevant flow environments. In particular, the impact of NA-mediated enzymatic desialylation on the characteristics of RBC aggregation, such as aggregate size, stability, and morphological structure, remains inadequately characterized. As a result, the specific contribution of glycocalyx degradation to RBC aggregation dynamics under flow conditions continues to be an open and insufficiently explored question.

To address the unresolved role of glycocalyx degradation in RBC aggregation under flow, we employed NA from *C. welchii*, a well-characterized and dose-controllable enzyme, to simulate varying levels of desialylation. In our previous studies, we have shown that treatment with ∼90 units/L (U/L) of NA removes most sialic acid residues from the RBC surface [25, 26], aligning with pathological activity levels of the enzyme observed in conditions such as diabetes (50—65 U/L) [23, 25, 26]. In our study, RBCs were treated with NA across a range of activity 0—100 U/L, followed by quantitative assessment of glycocalyx alterations via confocal microscopy and analysis of aggregation behavior under shear flow using microfluidic platform. While classical models of RBC aggregation, namely the depletion and bridging model [2, 27, 28, 29, 30], emphasize the role of macromolecules from plasma or artificial adjunction of dextran, we hypothesize that glycocalyx structures modulate aggregation dynamics by influencing intercellular charge interactions, molecular spacing, and hydration forces. Through this work, we aim to provide new insights into how enzymatic degradation of specific glycocalyx components influences RBC aggregation under physiological flow conditions. This has particular relevance to inflammatory and metabolic pathologies where glycocalyx degradation is prevalent and may offer a clearer mechanistic understanding of altered blood rheology in pathological states.

## 2 Materials and methods

### 2.1 Microfluidics

All microfluidic devices were fabricated using standard soft lithography. In brief, a master mold of the microchannel was obtained from a negative photoresist (SU8, Gersteltec, Switzerland) spin-coated onto a silicon wafer and exposed to UV light by the direct laser writing equipment (Dilase 250, KLOÉ, France). Polydimethylsiloxane (PDMS, Sylgard 184, Dow Corning, USA) was cast onto the mold and left to cure for 2 hours at 65 °C. A glass coverslips (#1, thickness 150 *µ*m) were used as the bottom part of the microchips. It was permanently sealed to the PDMS upper part after exposure the surfaces of both elements to an oxygen plasma (PDC-32G-2, Harrick, USA). The microfluidic systems employed in this study featured a parallelized network of several straight PDMS channels (length L=3 cm, width w=45 *µ*m, height h=10 *µ*m), symmetrically branching from a common inlet and outlet reservoir Figure 1. This biomimetic design replicates post-capillary bed geometry [31], while parallelization ensures experimental throughput and minimizes inter-device variability. Transfer tubing (0.020-inch ID × 0.060-inch OD, Masterflex®, Avantor, USA) interfaced the chip with a fluid reservoir at the inlet and outlet, ensuring leak-free operation up to 2000 mbar.

**Figure 1:**
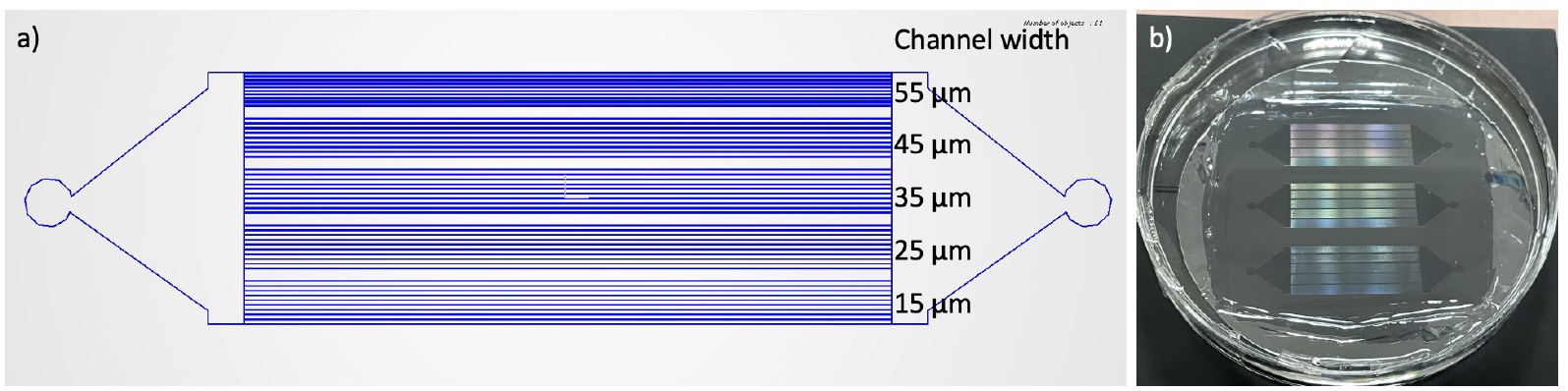
Design of microfluidic chip. a) Design of the straight microfluidic channels (the width is 15, 25, 35, 45, and 55 *µ*m). b) Overall view of wafer with printed circuit.

A pressure-driven flow was established using a microfluidic pressure controller (OB1 MK3, Elveflow, France) to pressurize the inlet reservoir containing RBC suspensions. The pressure drop (*△*P) between inlet and outlet was regulated within 30—150 mbar via the Elveflow Smart Interface software (ESI, Elveflow, France). For rectangular microchannels, the wall shear stress (*τ*) was derived from the force balance between pressure and viscous forces:

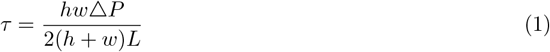

where L=3 cm, w=45 *µ*m, h=10 *µ*m. This yielded *τ*=0.40—2.05 Pa, encompassing physiological shear stress ranges in human microvasculature [31, 32, 33]. In subsequent experiments, discrete shear stress values of approximately 0.40, 0.68, 0.95, 1.23, 1.50, 1.77 and 2.05 Pa were applied to represent low to high physiological flow conditions.

### 2.2 Red blood cell samples

Whole blood samples were provided by the Établissement du Sang (EFS Grenoble, France) from healthy donors and stored in a citrated tube at 4 °C up to 3 days before use. Whole blood was centrifuged at 6000 rpm for 3 minutes, and the supernatant (plasma) and buffy coat were gently removed by aspiration. The pelleted RBCs were resuspended in phosphate buffer saline (PBS, Sigma-Aldrich, USA) solution and washed by aspiration and discharge, followed by centrifugation again. These washing steps were repeated 3 times to ensure that only RBCs remained in the suspension. Washed RBCs (considered as a 100% hematocrit suspension) would be resuspended in different solutions for various objectives (aggregation or enzymatic treatment). All experiments were repeated at least 5 times using samples from different healthy donors.

For glycocalyx modification, the washed RBCs at 10% hematocrit were exposed to NA from *Clostridium perfringens* (*C. welchii*, N2876, Sigma-Aldrich, USA) in PBS solution at different activity concentrations (10, 25, 50, 75, or 100 U/L) to disrupt glycocalyx. This incubation was performed on the roller (BioCote®, UK) at room temperature for 3 hours, keeping a homogenous solution and avoiding sedimentation of RBCs. The incubation was followed by 3 times washing in PBS. Unless otherwise indicated, untreated RBCs incubated under identical conditions without NA were used as the control.

For microfluidic experiments, washed RBCs were suspended at 5% hematocrit in a PBS solution supplemented with 15 mg/mL dextran from *Leuconostoc mesenteroides* (M_w_*≈*150 kDa, Sigma-Aldrich, USA).

For the fluorescence microscopy experiment, washed RBCs with or without NA treatment (5% hematocrit in PBS) were incubated with 5 *µ*g/mL Maackia Amurensis Lectin II (MAL II) conjugated to Texas Red (GlycoMatrix™, USA) for 1 hour at room temperature. Unbound lectin was removed by 3 cycles of centrifugation and resuspension in PBS solution, and labeled RBCs were resuspended to 5% hematocrit for imaging. Additionally, the autofluorescence of RBCs without NA treatment and without labeling with MAL/MAL II was verified as a negative control to ensure accurate measurements.

### 2.3 Image acquisition

Confocal fluorescence imaging of MAL II-stained RBCs was performed on an inverted microscope Leica SP8 (Leica Microsystems GmbH, Germany). XYZ stacks were obtained with 40×/NA1.30 oil-immersion objective in raster mode, with a lateral image size of 512 × 512 pixels, a lateral resolution of 0.263 *µ*m/pixel, and a Z-slice spacing of 0.35 *µ*m. All confocal imaging was carried out using identical acquisition settings across experimental groups to ensure comparability.

Unstained RBCs, under both static and flow conditions, were visualized in bright-field mode using an Olympus inverted microscope (Olympus Corporation, Japan) with a 455 nm LED light source, a 60×/NA 0.70 high-precision objective(Olympus Corporation, Japan), and a Photron Mini UX50 highspeed camera (Photron, Japan). 12-bit monochrome images with a resolution of 1280×1024 pixels (0.167 *µ*m/pixel) were captured at a rate of 2000 fps. Acquisition parameters were kept constant for all experimental conditions.

### 2.4 Image analysis and processing

Human blood samples from distinct donors were used to account for biological variability in the experimental design. For each concentration and/or each wall shear stress condition, a substantial number of images were captured to enable statistically robust quantification of transient aggregation phenomena. At least 5 independent replicates were performed for all experiments, with donor samples randomized across trials to mitigate potential batch effects. The acquired image stacks were processed and analyzed using the FIJI/ImageJ2 open-source platform and its built-in plugins to extract relevant parameters [34].

#### Fluorescence quantification (MAL II)

The confocal 3D image stacks with dark backgrounds were analyzed to evaluate glycocalyx modifications. For each sample, a minimum of 10 randomly selected fields of view were acquired under identical imaging conditions. To minimize bias, all Z-stacks were processed in their entirety rather than selecting individual slices. Background signal was removed using the “Subtract Background” function with a rolling-ball radius of 60 pixels to eliminate low-frequency noise while preserving the cell signal. For each Z-stack, integrated fluorescence intensity was calculated by using the “Measure Stack” function after background subtraction. To account for differences in acquisition depth and field-to-field variation, the resulting values were normalized using a min–max normalization procedure as follows:

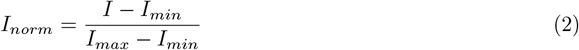

where I is the raw intensity, and I_min_ and I_max_ are the minimum and maximum intensities within the dataset. The number of RBCs in the same field was determined by manual counting on the corresponding projection, and the normalized fluorescence intensity per cell was calculated as the ratio of integrated I_norm_ to the RBC count. Representative values for each donor and treatment condition were obtained by averaging across all analyzed fields.

#### Bright-field aggregation analysis

For unstained RBC aggregation analysis, bright-field images were processed in the following order: (1) background subtraction with a rolling-ball filter (radius 60 pixels); (2) optional median filtering (2 pixels) to remove high-frequency noise; (3) global thresholding to generate binary masks; (4) morphological cleaning (opening/closing, 1–2 pixels) to remove artifacts; and (5) particle analysis using “Analyze Particles”. For each aggregate, the projected area (A) and perimeter (P) were measured, and the aggregate shape parameter circularity (ASP) was calculated as follows:

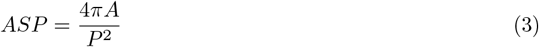

Here, ASP = 1 corresponds to a perfect circle, and ASP → 0 to increasingly elongated shapes. The number of RBCs per aggregate (N_agg_) was estimated as the ratio of aggregate area to the mean projected area of single RBCs measured from the same dataset. Local hematocrit was defined as the fraction of the total RBC area relative to the total image area. Particles smaller than 0.5 × the mean RBC area were excluded as debris.

For static aggregation assays, at least 20 images were analyzed per sample, and each condition was tested with RBCs from at least five independent healthy donors. For flow experiments, blood samples from healthy donors were analyzed for each NA concentration, and more than 20000 images per condition were processed. Aggregate size distributions were derived by classifying aggregates according to the number of RBCs they contained and calculating the relative frequency of each size class over the total number of aggregates. This probability distribution was then used to compare the occurrence of small (2–6 RBCs per aggregate), medium (7–11 RBCs per aggregate), and large aggregates (*≥*12 RBCs) across NA concentrations.

To ensure clarity, we distinguish between single-object and averaged parameters. N_agg_ refers to the number of RBCs in a single aggregate, whereas ⟨N_agg_⟩ denotes the average aggregate size across all aggregates within a sample. Similarly, ASP represents the circularity of an individual aggregate, while ⟨ASP⟩ is the mean circularity calculated across the aggregate population. Only aggregates containing more than 10 RBCs were included in the calculation of ⟨ASP⟩ to minimize bias from small near-spherical clusters.

#### Statistical analysis

To systematically evaluate the effects of different NA activities and flow conditions on RBC aggregation dynamics, human blood samples were analyzed as described above to account for biological variability. For each NA concentration and/or wall shear stress, a large number of images was acquired to enable statistically robust quantification of transient aggregation phenomena.

For each experimental condition, all individual cell-level measurements obtained across multiple donors were aggregated to characterize the overall distribution of the aggregation response. Sampling from several independent donors increased biological diversity and ensured that observed trends were not driven by a single individual. Mean values and associated variability were computed from the pooled measurements to describe the global phenotype under each condition.

Statistical comparisons between groups were performed using one-way analysis of variance (ANOVA, *α*=0.05) followed by Tukey’s post-hoc test for pairwise comparisons (GraphPad Prism 9.0, USA). All quantitative data are presented as mean±SEM, and differences were considered statistically significant at *p<*0.05. This analytical strategy allowed robust evaluation of intergroup differences without assuming donor-level dependence for cell-resolved measurements, thereby enhancing the reliability of the reported results.

## 3 Results

### 3.1 RBC surface desialylation induced by neuraminidase

The representative fluorescence microscopy images and quantitative results are presented in Figure 2. A direct comparison between the MAL II-labeled 0 U/L control group and RBCs treated with NA reveals a substantial reduction in fluorescence intensity in the presence of the enzyme, indicating progressive removal of sialic acid residues from the RBC surface. As shown in Figure 2, the normalized fluorescence intensity of the RBC glycocalyx decreased in a dose-dependent manner with increasing NA concentration (*p<*0.05 compared with control). Overall, fluorescence intensity exhibited a downward trend across the 0–100 U/L range, although the magnitude of the decrease varied among groups. These results confirm that NA effectively cleaves the MAL II-binding sialylated glycans on the RBC membrane.

**Figure 2:**
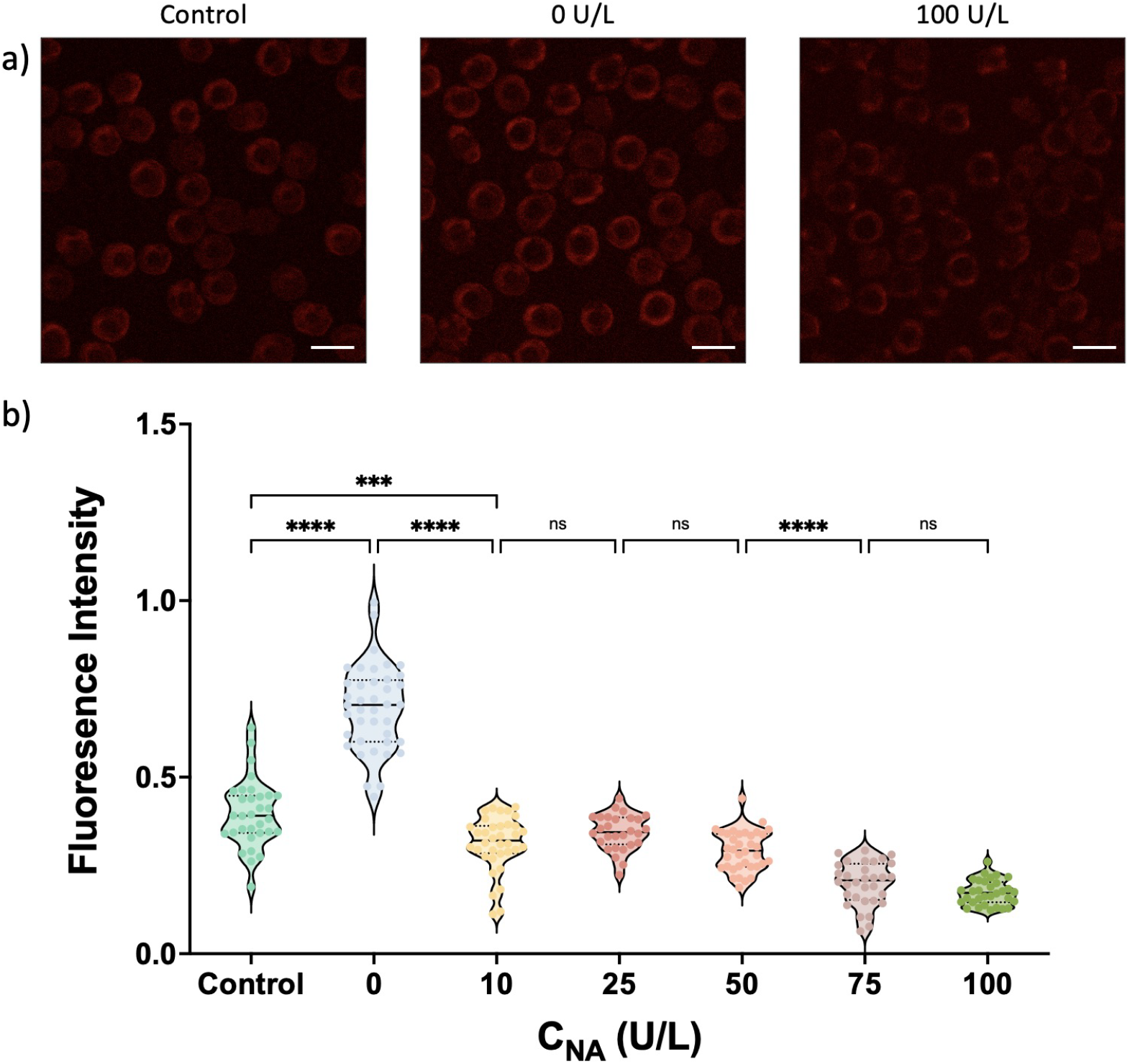
Staining of RBC glycocalyx sialic acids using Texas Red-conjugated MAA/MAL II (scale bar: 10 *µ*m). a) Representative confocal image of RBCs under three conditions: negative control, untreated, and neuraminidase-treated (100 U/L). b) Quantification of normalized fluorescence intensity as a function of NA activity (\*\*\**p <* 0.001, \*\*\*\**p <* 0.0001).

### 3.2 Impact of glycocalyx desialylation on RBC aggregation in static conditions

We first evaluated the effect of NA-induced glycocalyx degradation on RBC aggregation under static conditions. Dextran (M_w_*≈*150 kDa) was used to promote aggregation in PBS. RBC aggregation was quantified using average N_agg_ as a measure of aggregate size and average ASP as a measure of aggregate morphology, as defined in Equation 3.

Figure 3 shows representative images of RBC aggregates treated with various activities of NA. Qualitative observations indicate that enzymatic cleavage of the glycocalyx promotes the formation of larger and more elongated aggregates, with some samples showing highly branched or irregular morphologies.

**Figure 3:**
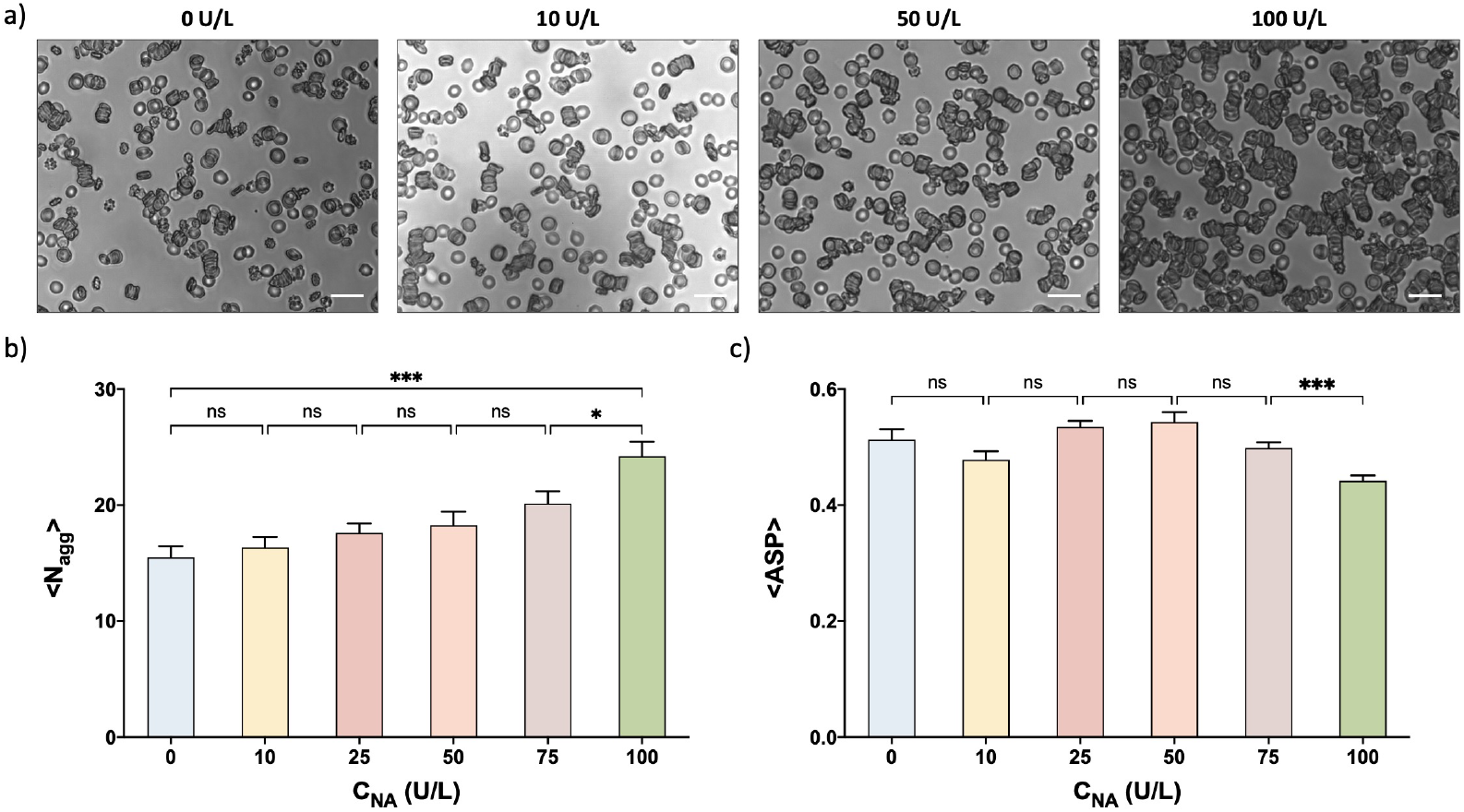
Static aggregation of RBCs following NA treatment (scale bar: 20 *µ*m). a) Representative microscopic images of RBC aggregates in static conditions after treatment with different NA activity levels. b) Average number of RBCs per aggregate (mean±SEM) under static conditions for each NA activity level (\**p <* 0.05, \*\*\**p <* 0.001). c) Average aggregate circularity (mean±SEM) in static conditions following treatment with different NA activity levels (\*\*\**p <* 0.001).

As shown in Figure 3, ⟨N_agg_⟩, the mean of N_agg_, increased with NA activity. No statistically significant difference was detected between the control group and samples treated with 10, 25, 50 or 75 U/L NA. In contrast, treatment with 100 U/L NA significantly increased aggregate size compared with the untreated control. These results indicate that NA treatment promotes RBC aggregation under static conditions.

To characterize aggregate morphology, we quantified the shape parameter ASP defined by Equation 3, which reflects the two dimensional circularity of aggregates. To reduce the influence of small aggregates that may appear nearly spherical, only aggregates containing more than 10 RBCs were included in this analysis.

As shown in Figure 3, ⟨ASP⟩ did not exhibit a clear dose-dependent trend with increasing NA activity. Significant differences were observed only between samples treated with 75 and 100 U/L NA, whereas no significant differences were detected for the other treatment groups. Representative images nevertheless showed that NA-treated samples contained larger aggregates with more irregular and branched morphologies. Together, these results suggest that under static conditions, ASP alone does not fully capture the morphological changes induced by NA treatment.

### 3.3 Stability of RBC aggregates in a straight microchannel in flow conditions

#### 3.3.1 Influence of glycocalyx degradation on RBC aggregate formation in flow conditions

To investigate the impact of sialic acid residue removal in the RBC glycocalyx on aggregate formation under flow, we conducted microfluidic experiments across varying NA activity levels. A range of pressure differences (*△*P=30–150 mbar) was applied to generate wall shear stresses spanning 0.40–2.05 Pa, corresponding to physiological conditions observed in microcirculation. Representative images under different treatment levels are shown in Figure 4.

**Figure 4:**
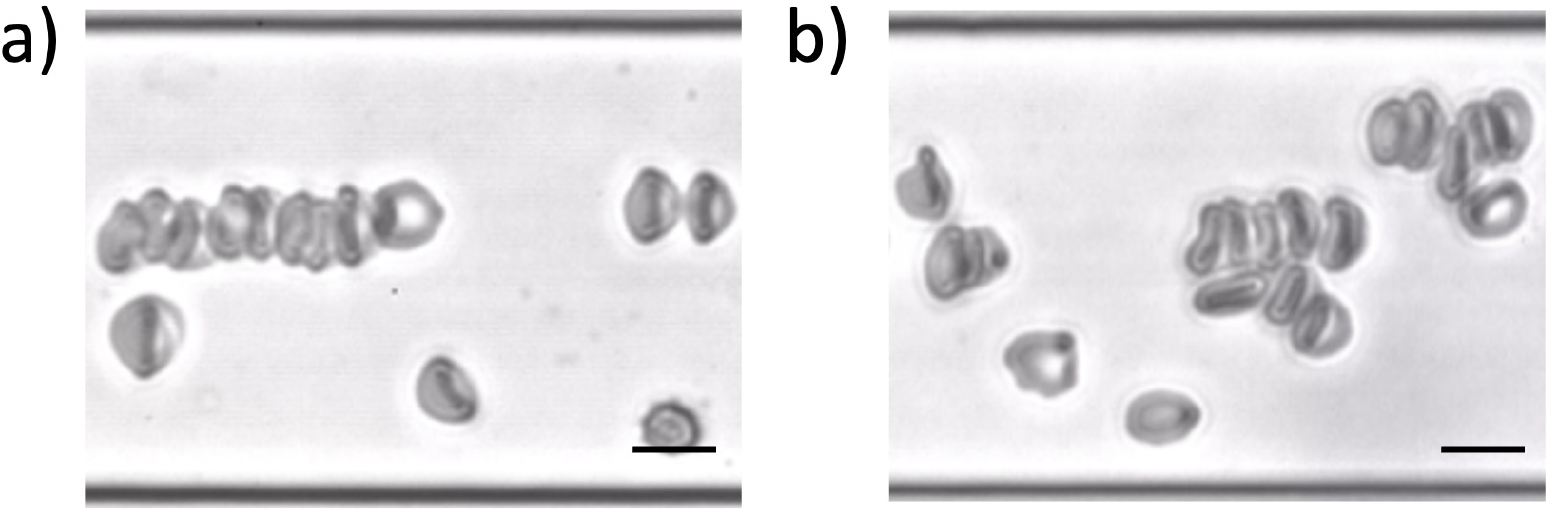
Representative microscopic images of RBC aggregates in a 45*µ*m wide microchannel under flow (*τ*=0.4 Pa; flow direction: left to right; scale bar: 10 *µ*m). a) Untreated RBCs (C_NA_=0 U/L). b) RBCs treated with NA (C_NA_=100 U/L).

The aggregate size distribution was first visualized by plotting the probability of different aggregate sizes occurring (Figure 5). This representation provides a general impression of aggregation behavior, however, the large variety of aggregate sizes complicates group-wise comparisons. Nonetheless, as shown in Figure 5, NA treatment (especially at 100 U/L) markedly increased the occurrence of large aggregates while reducing the frequency of small aggregates, suggesting that sialic acid cleavage facilitates the formation of larger clusters under flow. Similar trends were observed across the range of shear stresses investigated.

**Figure 5:**
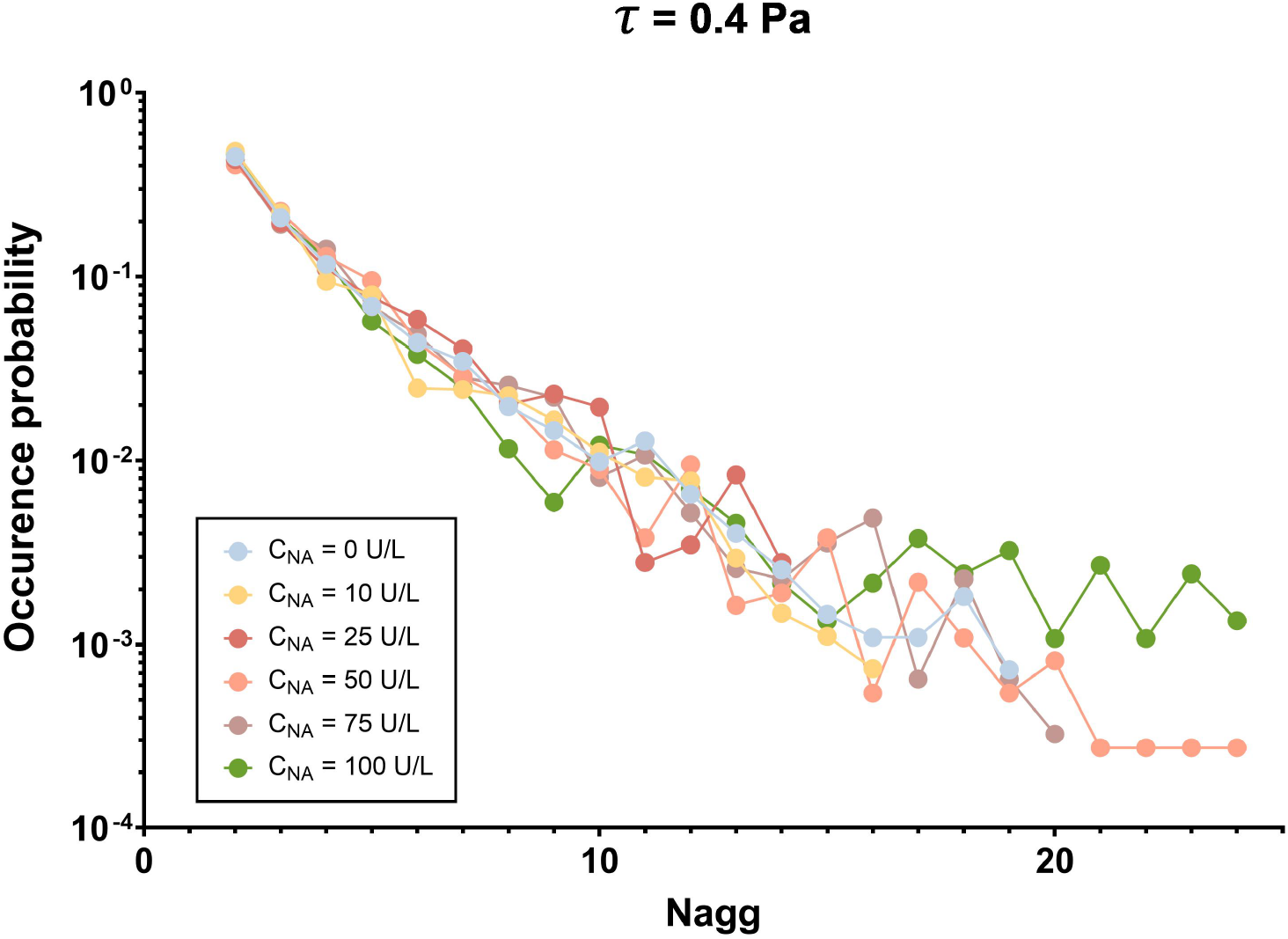
Size distribution of the RBC aggregates under different NA activity levels in a 45 *µ*m wide microchannel at shear stress *τ*=0.40 Pa. The 0.40 Pa condition is shown as a representative example.

Consistent with this observation, the largest aggregates detected in the experiments contained substantially more RBCs at elevated NA activities. While the exact values varied between samples, aggregates containing up to approximately 25–30 RBCs were observed in the highest NA groups, compared with approximately 15–20 RBCs in untreated controls.

Figure 6 further illustrates the dependence of the average aggregate size ⟨N_agg_⟩ on wall shear stresses (0.40–2.05 Pa) for different NA activity levels (0–100 U/L). To minimize the influence of hematocrit (Ht) variations across experiments, all values were normalized using the Ht-adjusted function (1+0.117×Ht), as derived from simulations in our previous work [35].

**Figure 6:**
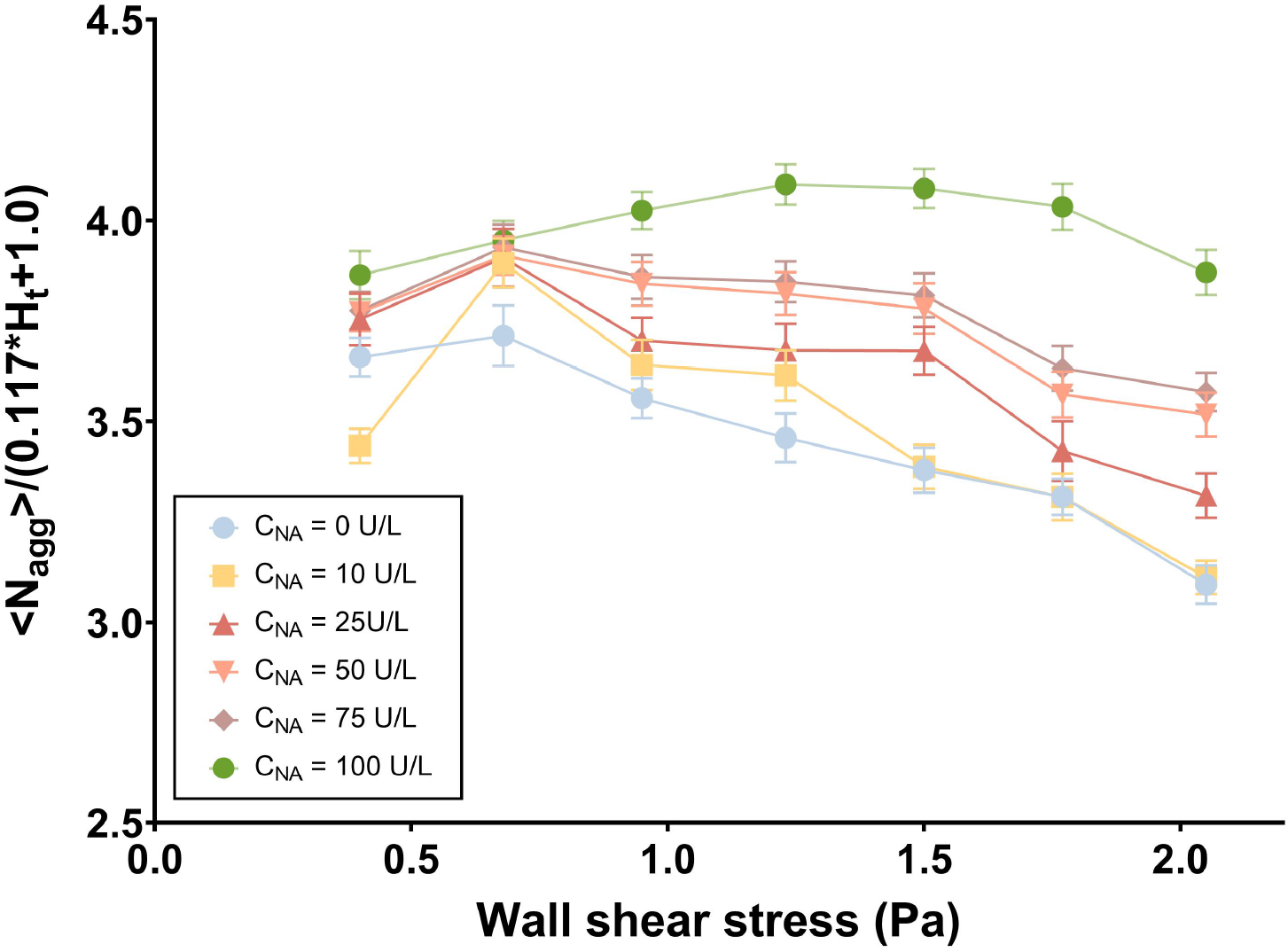
Normalized average size of RBC aggregates (mean±SEM) across NA concentrations by using a Ht-adjusted function [35].

At 0.40 Pa, aggregates were relatively small but stable. As shear increased to 0.68 Pa, ⟨N_agg_⟩ reached its maximum in the control and all groups treated with up to 75 U/L NA, showing that moderate shear promotes compaction or rearrangement. Beyond this point, ⟨N_agg_⟩ progressively decreased from 0.95 to 1.23 Pa, reflecting the onset of shear-induced disaggregation.

A different behavior was observed in the C_NA_=100 U/L group. Rather than reaching a maximum at 0.68 Pa, ⟨N_agg_⟩ continued to increase and peaked at 1.23 Pa before declining at higher stress. Even at 2.05 Pa, aggregates in the C_NA_=100 U/L condition remained larger than those in all other groups.

These results indicate that NA-induced desialylation modifies the shear-dependent behavior of RBC aggregates. While most groups followed a similar trend of initial growth (0.40–0.68 Pa) followed by progressive disaggregation (0.95–2.05 Pa), the highest NA activity shifted this transition toward higher shear stresses, resulting in aggregates that persisted over a broader mechanical range.

Collectively, these findings demonstrate that enzymatic removal of sialic acid residues from the RBC surface modulates aggregation behavior in a dose-dependent manner. NA treatment enhances intercellular adhesion by reducing electrostatic repulsion, resulting in the formation of larger and more shear-resistant RBC aggregates. This altered aggregation profile reflects a dynamic interplay between biochemical modifications of the glycocalyx and mechanical shear forces in the microenvironment. As a result, desialylated RBCs exhibit increased aggregate stability even under elevated wall shear stress, highlighting a potential mechanism by which glycocalyx degradation contributes to impaired microcirculatory flow. These effects may be particularly relevant in pathological conditions such as inflammation, diabetes, or sepsis, where compromised glycocalyx integrity may exacerbate vascular resistance and promote microvascular obstruction [36, 37, 38, 39, 40, 41, 42, 43].

#### 3.3.2 Morphology and structural stability of RBC aggregates in flow conditions

As previously observed in static conditions Figure 3, enzymatic digestion of the glycocalyx altered the shape of RBC aggregates while ⟨ASP⟩ values remain relatively stable. Under flow, however, the mechanical environment becomes a key determinant of aggregate morphology. As shown in Figure 4, compared to the typical rouleaux structures formed by untreated RBCs under physiological conditions (C_NA_=0 U/L), aggregates formed by NA-treated RBCs display not only size differences but also noticeable changes in shape. To further quantify this morphological change, we analyzed the relationship between average aggregate circularity (⟨ASP⟩) and size (N_agg_) across shear stresses ranging from 0.40 to 2.05 Pa and for different NA concentrations, as illustrated in Table1 and Figure S1.

**Table 1:** Representative values of ⟨ASP⟩ (mean±SEM) at selected aggregate sizes (N_agg_) and shear stresses (0.40 and 2.05 Pa) for different NA concentrations.

| $\tau$ | $C_{\text{NA}}$ (U/L) | $\langle \text{ASP} \rangle$ | | | |
| --- | --- | --- | --- | --- | --- |
| | | $N_{\text{agg}}=6$ | $N_{\text{agg}}=8$ | $N_{\text{agg}}=10$ | $N_{\text{agg}}=12$ |
| 0.40 Pa | 0 | $0.315 \pm 0.009$ | $0.265 \pm 0.010$ | $0.214 \pm 0.011$ | $0.188 \pm 0.011$ |
| | 25 | $0.332 \pm 0.007$ | $0.283 \pm 0.012$ | $0.290 \pm 0.019$ | $0.172 \pm 0.010$ |
| | 100 | $0.348 \pm 0.006$ | $0.268 \pm 0.010$ | $0.284 \pm 0.008$ | $0.274 \pm 0.028$ |
| 2.05 Pa | 0 | $0.274 \pm 0.007$ | $0.236 \pm 0.009$ | $0.200 \pm 0.012$ | $0.168 \pm 0.020$ |
| | 25 | $0.301 \pm 0.008$ | $0.234 \pm 0.010$ | $0.243 \pm 0.019$ | $0.187 \pm 0.015$ |
| | 100 | $0.308 \pm 0.005$ | $0.243 \pm 0.007$ | $0.219 \pm 0.007$ | $0.192 \pm 0.010$ |

Across all groups, a consistent trend was observed: larger aggregates were associated with lower ⟨ASP⟩ values, indicating more elongated or anisotropic structures. For example, in the control group (C_NA_=0 U/L), ⟨ASP⟩ decreased from ∼0.315 at N_agg_=6 to ∼0.188 at N_agg_=12 under 0.40 Pa. At higher shear stress (2.05 Pa), the corresponding values declined further to ∼0.274 and ∼0.168, respectively, reflecting stronger deformation.

Comparison across treatment groups revealed subtle but systematic differences. At N_agg_=6 and *τ*=0.40 Pa, the average ⟨ASP⟩ values were ∼0.315 (C_NA_=0 U/L), ∼0.332 (C_NA_=25 U/L), and ∼0.348 (C_NA_=100 U/L). Similar differences persisted at *τ*=2.05 Pa, with NA-treated samples consistently showing slightly higher ⟨ASP⟩ than controls at the same aggregate size and the largest differences observed in the C_NA_=100 U/L group.

Together, these findings indicate that aggregate morphology is influenced by both aggregate size and shear stress. Larger aggregates generally exhibited lower ASP values, while NA-treated samples maintained slightly higher ⟨ASP⟩ values than controls at matched aggregate sizes, particularly at the highest NA activity level.

## 4 Discussion and conclusion

This study demonstrates that enzymatic degradation of the RBC glycocalyx by NA significantly alters RBC aggregation behavior under physiologically relevant shear flow. Progressive removal of surface sialic acids increased aggregate size and promoted the persistence of aggregates under shear stress. While the effects were modest at low NA activities, higher enzyme concentrations produced larger aggregates that remained stable across a broad range of wall shear stresses. These findings support the concept that glycocalyx integrity is a critical determinant of RBC-RBC interactions and contributes directly to the regulation of blood rheology.

The role of sialic acids in RBC aggregation has been recognized for several decades. Sialic acid residues are predominantly carried by glycophorin A and related membrane glycoproteins, which account for a large fraction of the negative surface charge of RBCs. This negative charge generates electrostatic repulsion between neighboring cells and thereby limits spontaneous aggregation. Loss of membrane sialylation has been associated with erythrocyte aging, inflammation, infection, diabetes, and other pathological conditions, where excessive desialylation contributes to altered hemorheological properties and accelerated RBC clearance [44, 43, 45]. Consistent with these observations, our MAL II fluorescence measurements demonstrated a progressive loss of surface sialic acids following NA treatment, accompanied by a corresponding increase in RBC aggregation. Together, these findings provide direct experimental evidence linking glycocalyx desialylation to enhanced aggregation under controlled flow conditions.

Most previous studies investigating the relationship between RBC desialylation and aggregation have relied on static aggregation assays or bulk rheological measurements. While these approaches have provided valuable insights into the role of surface sialic acids in regulating cell-cell interactions, they do not fully capture the dynamic mechanical environment experienced by RBCs in the microcirculation. By combining controlled NA-mediated desialylation with microfluidic flow assays, the present study extends these observations to physiologically relevant shear conditions. This approach enables direct quantification of aggregate formation, morphology, and persistence under flow, providing a more comprehensive assessment of how glycocalyx degradation may influence microvascular blood transport.

Beyond aggregate size, our results suggest that desialylation may also influence aggregate morphology. Under flow conditions, larger aggregates exhibited lower ASP values, reflecting increasingly elongated or anisotropic structures. In addition, NA-treated RBCs consistently maintained slightly higher ASP values than untreated controls at matched aggregate sizes, particularly at the highest enzyme concentrations. Although the present study does not directly resolve the internal architecture of aggregates, these observations suggest that desialylated RBCs may organize differently during aggregate formation. Rather than forming exclusively classical rouleaux structures, desialylated RBCs may adopt more heterogeneous aggregate configurations under flow. Future high-resolution imaging studies will be required to determine whether glycocalyx degradation alters aggregate packing and structural organization.

Beyond its role in regulating surface charge, membrane sialylation may also influence the structural organization of the erythrocyte membrane. Sialic acid residues are primarily carried by glycophorins, which are closely associated with membrane proteins linked to the underlying cytoskeleton. The RBC membrane is mechanically supported by a spectrin-actin cytoskeletal network connected to membrane proteins such as Band 3, ankyrin, and glycophorin A [46, 47]. Previous studies have suggested that membrane glycosylation and sialylation contribute not only to surface charge but also to membrane organization, protein mobility, and cellular mechanical properties [44, 48]. Although the present study did not directly investigate membrane-cytoskeleton interactions, it is conceivable that extensive desialylation may influence the coupling between the glycocalyx, plasma membrane, and cytoskeletal network. Such effects could potentially contribute to changes in RBC deformability, aggregation behavior, and mechanical stability. Future studies integrating glycocalyx characterization, membrane mechanics, and cytoskeletal analyses will be required to evaluate this possibility.

The pathological relevance of these findings extends beyond RBC-RBC interactions. Elevated NA activity has been reported in several diseases, including influenza, bacterial infections, diabetes, and sepsis, all of which are associated with vascular complications and impaired microcirculation [11, 23, 42, 43, 45]. Increased RBC aggregation has long been recognized as an important contributor to elevated blood viscosity and flow heterogeneity in these conditions [40, 37]. Our results therefore provide a mechanistic framework linking NA-mediated desialylation to altered RBC rheology. By promoting the formation of larger and more persistent aggregates, glycocalyx degradation may contribute to impaired perfusion and increased vascular resistance within the microcirculation.

Importantly, glycocalyx degradation is unlikely to affect only RBCs *in vivo*. Endothelial cells also possess a glycocalyx-rich surface layer that plays a fundamental role in vascular homeostasis. Consequently, NA activity may simultaneously alter RBC-RBC and RBC-endothelium interactions. To investigate such interactions under physiologically relevant conditions, endothelialized microfluidic platforms have been developed to reproduce key features of the microcirculation [49, 50]. Using a similar experimental approach, our preliminary observations suggest that glycocalyx degradation increases RBC residence time on endothelial surfaces, particularly at vascular bifurcations where local flow disturbances occur. Although these findings remain preliminary, they emphasize the importance of considering both erythrocyte and endothelial glycocalyx integrity when evaluating the microvascular consequences of desialylation.

Taken together, our findings identify glycocalyx desialylation as an important regulator of RBC aggregation under flow conditions. By linking NA activity, loss of surface sialic acids, aggregate formation, and altered aggregate morphology, this study provides new insight into the mechanisms governing blood cell interactions in the microcirculation. Future investigations integrating glycocalyx biology, membrane mechanics, membrane-cytoskeleton coupling, and RBC-endothelium interactions will be essential for understanding how desialylation contributes to microvascular dysfunction in pathological states.

## 5 Supplementary Materials

**Figure S1:**
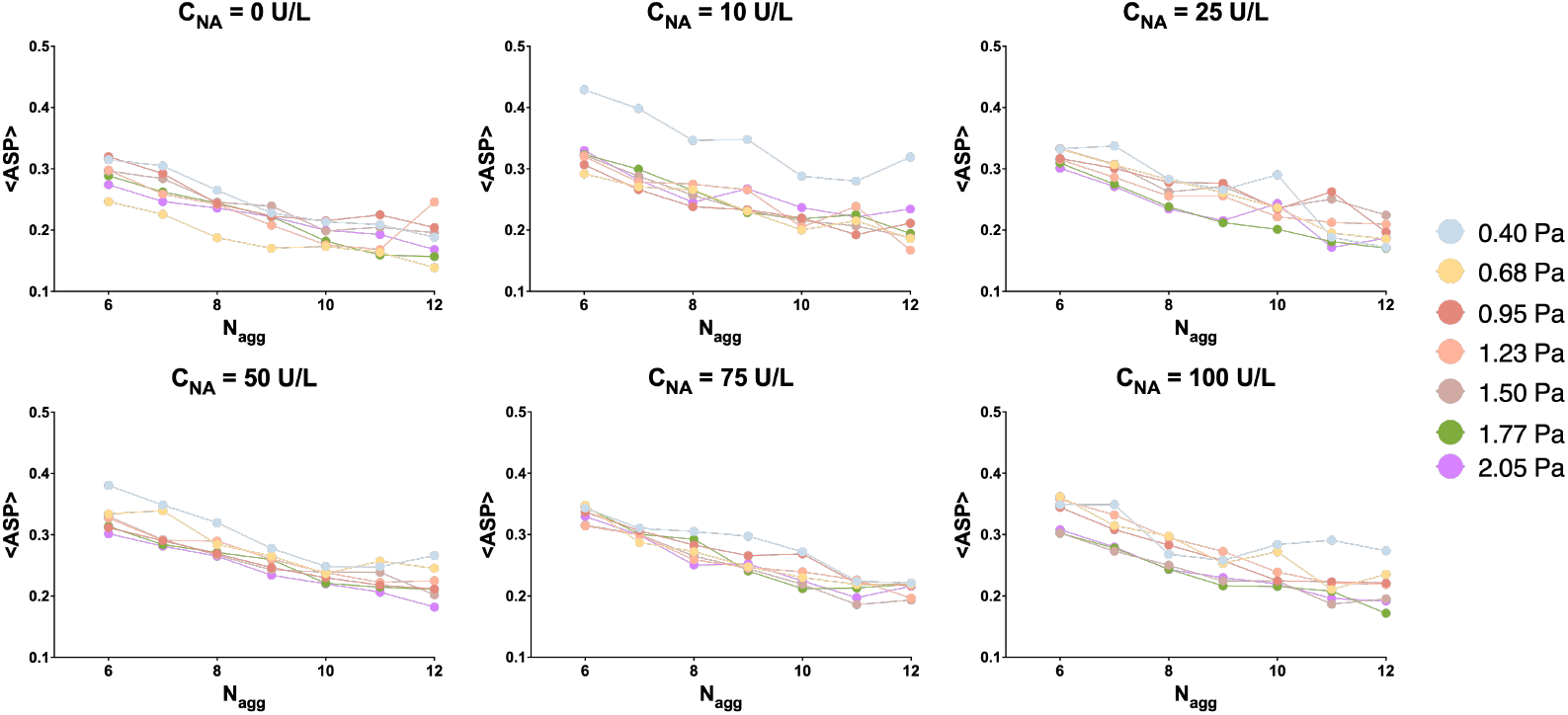
Average shape parameter (⟨ASP⟩) as a function of the aggregate size (N_agg_) under the flow with different wall shear stress.

## References

[1] O. K. Baskurt and H. J. Meiselman. Erythrocyte aggregation: Basic aspects and clinical importance. Clinical Hemorheology and Microcirculation, 53(1-2):23–37, 2013.

[2] M. Brust, O. Aouane, M. Thiébaud, D. Flormann, C. Verdier, L. Kaestner, M. W. Laschke, A. Selmi, H.and Benyoussef, T. Podgorski, G. Coupier, C. Misbah, and C. Wagner. The plasma protein fibrinogen stabilizes clusters of red blood cells in microcapillary flows. Scientific Reports, 4:4348, 2014.

[3] A. S. Popel and P. C. Johnson. Microcirculation and hemorheology. Annual Review of Fluid Mechanics, 37:43–69, 2005.

[4] V. D. O’Loughlin, E. Pennefather-O’Brien, and M. P. McKinley. Human Anatomy. McGraw Hill LLC, New York, 2024.

[5] E. H. Eylar, M. A. Madoff, O. V. Brody, and J. L. Oncley. The contribution of sialic acid to the surface charge of the erythrocyte. Journal of Biological Chemistry, 237:1992–2000, 1962.

[6] G. M. W. Cook, D. H. Heard, and G. V. F. Seaman. Sialic acids and the electrokinetic charge of the human erythrocyte. Nature, 191:44–47, 1961.

[7] O. K. Baskurt. Handbook of Hemorheology and Hemodynamics. IOS Press, Amsterdam, 2007.

[8] S. Chien, S. Usami, H. M. Taylor, J. L. Lundberg, and M. I. Gregersen. Effects of hematocrit and plasma proteins on human blood rheology at low shear rates. Journal of Applied Physiology, 21(1):81–87, 1966.

[9] S. Chien, S. Usami, R. J. Dellenback, M. I. Gregersen, L. B. Nanninga, and M. M. Guest. Blood viscosity: Influence of erythrocyte aggregation. Science, 157(3790):829–831, 1967.

[10] S. Chen, A. Eldor, G. Barshtein, S. Zhang, A. Goldfarb, E. Rachmilewitz, and S. Yedgar. Enhanced aggregability of red blood cells of beta-thalassemia major patients. The American Journal of Physiology, 270(6 Pt 2):H1951–H1956, 1996.

[11] M. E. Rogers, D. T. Williams, R. Niththyananthan, M. W. Rampling, K. E. Heslop, and D. G. Johnston. Decrease in erythrocyte glycophorin sialic acid content is associated with increased erythrocyte aggregation in human diabetes. Clinical Science (London, England : 1979), 82(3):309–313, 1992.

[12] R. Kotan, K. Peto, A. Deak, Z. Szentkereszty, and N. Nemeth. Hemorheological and microcirculatory relations of acute pancreatitis. Metabolites, 13(1):4, 2022.

[13] E. J. Dunn and R. A. Ariëns. Fibrinogen and fibrin clot structure in diabetes. Herz, 29(5):470–479, 2004.

[14] G. Fusman, T. Mardi, D. Justo, M. Rozenblat, R. Rotstein, D. Zeltser, A. Rubinstein, M. Koffler, E. Shabtai, S. Berliner, and I. Shapira. Red blood cell adhesiveness/aggregation, c-reactive protein, fibrinogen, and erythrocyte sedimentation rate in healthy adults and in those with atherosclerotic risk factors. The American Journal of Cardiology, 90(5):561–563, 2002.

[15] S. Shin, Y. Ku, N. Babu, and M. Singh. Erythrocyte deformability and its variation in diabetes mellitus. Indian Journal of Experimental Biology, 45(1):121–128, 2007.

[16] G. Gambaro, B. Baggio, E. Cicerello, S. Mastrosimone, G. Marzaro, A. Borsatti, and G. Crepaldi. Abnormal erythrocyte charge in diabetes mellitus: Link with microalbuminuria. Diabetes, 37(6):745–748, 1988.

[17] J. Stuart and I. Juhan-Vague. Erythrocyte rheology in diabetes mellitus. Clinical Hemorheology and Microcirculation, 7(2), 1987.

[18] R. Agrawal, T. Smart, C. Richards, R. Bhatnagar, A. Tufail, D. Shima, and C. Pavesio. Assessment of red blood cell deformability in type 2 diabetes mellitus and diabetic retinopathy by dual optical tweezers stretching technique. Scientific Reports, 6:15873, 2016.

[19] J. A. Miller, E. Gravallese, and H. F. Bunn. Nonenzymatic glycosylation of erythrocyte membrane proteins: Relevance to diabetes. The Journal of Clinical Investigation, 65(4):896–901, 1980.

[20] J. Tripette, Allard L. Nguyen, L. C., P. Robillard, G. Soulez, and G. Cloutier. In vivo venous assessment of red blood cell aggregate sizes in diabetic patients with a quantitative cellular ultrasound imaging method: Proof of concept. PLoS ONE, 10(4):e0124712, 2015.

[21] W. Zhu, Y. Zhou, L. Guo, and S. Feng. Biological function of sialic acid and sialylation in human health and disease. Cell death discovery, 10(1):415, 2024.

[22] V. Escuret and O. Terrier. Co-infection of the respiratory epithelium, scene of complex functional interactions between viral, bacterial, and human neuraminidases. Frontiers in Microbiology, 14:1137336, 2023.

[23] A. Merat, R. Arabsolghar, J. Zamani, and M. H. Roozitalab. Serum levels of sialic acid and neuraminidase activity in cardiovascular, diabetic and diabetic retinopathy patients. Iranian Journal of Medical Sciences, 28(3):123–126, 2015.

[24] S. M. Qadri, D. A. Donkor, I. Nazy, D. R. Branch, and W. P. Sheffield. Bacterial neuraminidase-mediated erythrocyte desialylation provokes cell surface aminophospholipid exposure. European Journal of Haematology, 100(5):502–510, 2018.

[25] Y. Izumida, A. Seiyama, and N. Maeda. Erythrocyte aggregation: Bridging by macromolecules and electrostatic repulsion by sialic acid. Biochimica et Biophysica Acta, 1067(2):221–226, 1991.

[26] N. Maeda, K. Imaizumi, M. Sekiya, and T. Shiga. Rheological characteristics of desialylated erythrocytes in relation to fibrinogen-induced aggregation. Biochimica et Biophysica Acta, 776(1):151–158, 1984.

[27] J. K. Armstrong, R. B. Wenby, H. J. Meiselman, and T. C. Fisher. The hydrodynamic radii of macromolecules and their effect on red blood cell aggregation. Biophysical Journal, 87(6):4259–4270, 2004.

[28] N. Maeda, M. Seike, S. Kume, T. Takaku, and T. Shiga. Fibrinogen-induced erythrocyte aggregation: erythrocyte-binding site in the fibrinogen molecule. Biochimica et Biophysica Acta (BBA) - Biomembranes, 904(1):81–91, 1987.

[29] P. Steffen, C. Verdier, and C. Wagner. Quantification of depletion-induced adhesion of red blood cells. Physical Review Letters, 110(1):018102, 2013.

[30] B. Neu and H. J. Meiselman. Depletion-mediated red blood cell aggregation in polymer solutions. Biophysical Journal, 83(5):2482–2490, 2002.

[31] A. G. Koutsiaris, S. V. Tachmitzi, N. Batis, M. G. Kotoula, C. H. Karabatsas, E. Tsironi, and D. Z. Chatzoulis. Volume flow and wall shear stress quantification in the human conjunctival capillaries and post-capillary venules in vivo. Biorheology, 44(5–6):375–386, 2007.

[32] N. Baeyens, C. Bandyopadhyay, B. G. Coon, S. Yun, and M. A. Schwartz. Endothelial fluid shear stress sensing in vascular health and disease. Journal of Clinical Investigation, 126(3):821–828, 2016.

[33] S. Chatterjee. Endothelial mechanotransduction, redox signaling and the regulation of vascular inflammatory pathways. Frontiers in Physiology, 9:524, 2018.

[34] J. Schindelin, I. Arganda-Carreras, E. Frise, V. Kaynig, M. Longair, T. Pietzsch, S. Preibisch, C. Rueden, S. Saalfeld, B. Schmid, J. Y. Tinevez, D. J. White, V. Hartenstein, K. Eliceiri, P. Tomancak, and A. Cardona. Fiji: an open-source platform for biological-image analysis. Nature Methods, 9(7):676–682, 2012.

[35] M. Abbasi, M. Jin, Rashidi Y., L. Bureau, D. Tsvirkun, and C. Misbah. Glycocalyx cleavage boosts erythrocytes aggregation. Scientific Reports, 14:24340, 2024.

[36] A. E. Lugovtsov, Y. I. Gurfinkel, P. B. Ermolinskiy, A. I. Maslyanitsina, L. I. Dyachuk, and A. V. Priezzhev. Optical assessment of alterations of microrheologic and microcirculation parameters in cardiovascular diseases. Biomedical Optics Express, 10(8):3974–3986, 2019.

[37] D. K. Kaul. Sickle red cell adhesion: Many issues and some answers. Transfusion Clinique et Biologique: Journal de la Societe Francaise de Transfusion Sanguine, 15(1–2):51–55, 2008.

[38] A. Semenov, A. Lugovtsov, P. Ermolinskiy, K. Lee, and A. Priezzhev. Problems of red blood cell aggregation and deformation assessed by laser tweezers, diffuse light scattering and laser diffractometry. Photonics, 9(4):238, 2022.

[39] M. Levi, M. Schultz, and T. van der Poll. Sepsis and thrombosis. Seminars in Thrombosis and Hemostasis, 39(5):559–566, 2013.

[40] O. K. Baskurt, A. Temiz, and H. J. Meiselman. Red blood cell aggregation in experimental sepsis. The Journal of Laboratory and Clinical Medicine, 130(2):183–190, 1997.

[41] Y. S. Chang, C. H. Ho, C. C. Chu, J. J. Wang, and R. L. Jan. Risk of retinal vein occlusion in patients with diabetes mellitus: A retrospective cohort study. Diabetes Research and Clinical Practice, 171:108607, 2021.

[42] S. Ghosh. Sialic Acids and Sialoglycoconjugates in the Biology of Life, Health and Disease. Academic Press, London, 2020.

[43] B. Venerando, A. Fiorilli, G. Croci, C. Tringali, G. Goi, L. Mazzanti, G. Curatola, G. Segalini, L. Massaccesi, A. Lombardo, and G. Tettamanti. Acidic and neutral sialidase in the erythrocyte membrane of type 2 diabetic patients. Blood, 99(3):1064–1070, 2002.

[44] A. Varki. Biological roles of glycans. Glycobiology, 27(1):3–49, 2017.

[45] M. Matrosovich, T. Y. Matrosovich, T. Gray, N. A. Roberts, and H. D. Klenk. Neuraminidase is important for the initiation of influenza virus infection in human airway epithelium. Journal of Virology, 78(22):12665–12667, 2004.

[46] N. Mohandas and E. Evans. Mechanical properties of the red cell membrane in relation to molecular structure and genetic defects. Annual review of biophysics and biomolecular structure, 23:787–818, 1994.

[47] N. Mohandas and P. G. Gallagher. Red cell membrane: past, present, and future. Blood, 112(10):3939–3948, 2008.

[48] R. Shurer, J. C. Kuo, L. M. Roberts, J. G. Gandhi, M. J. Colville, T. A. Enoki, H. Y. Pan, J. Su, J. M. Noble, M. J. Hollander, J. P. O’Donnell, R. Yin, K. Pedram, L. Möckl, L. F. Kourkoutis,W. E. Moerner, C. R. Bertozzi, G. W. Feigenson, H. L. Reesink, and M. J. Paszek. Physical principles of membrane shape regulation by the glycocalyx. Cell, 177(7):1757–1770.e21, 2019.

[49] D. Tsvirkun, A. Grichine, A. Duperray, C. Misbah, and L. Bureau. Microvasculature on a chip: study of the endothelial surface layer and the flow structure of red blood cells. Scientific Reports, 7:45036, 2017.

[50] M. Inglebert, L. Locatelli, D. Tsvirkun, P. Sinha, J. A. Maier, C. Misbah, and L. Bureau. The effect of shear stress reduction on endothelial cells: A microfluidic study of the actin cytoskeleton. Biomicrofluidics, 14(2):024115, 2020.

